# Desmoplakin tail domain position in the desmosomal plaque is isoform dependent

**DOI:** 10.1101/2025.02.03.636241

**Authors:** Collin M. Ainslie, Krishna Patel, Yen T. B. Tran, Navaneetha Krishnan Bharathan, Volker Spindler, Alexa L. Mattheyses

## Abstract

Desmoplakin (DP) is the anchoring subunit of desmosomes, macromolecular junctions that provide mechanical integrity to the skin and heart. DP has three isoforms, DPI, DPIa, and DPII that arise from alternative splicing. The isoforms are structurally identical excluding the length of their central rod domain. As desmosomes are macromolecular complexes, the precise arrangement of their component proteins, or architecture, is essential to maintain physiological function. Alterations of the tissue-specific expression of DP isoforms underlies rare human diseases impacting the skin and heart. Overall DP is oriented with its head domain closest to the plasma membrane and tail domain extending into the cytosol. However, the differences in the architecture of the DP isoforms within the desmosomal plaque remains unknown. Here, we sought to define the architectural arrangement of each DP isoform. To address this, we utilized direct stochastic optical reconstruction microscopy (dSTORM) and analysis of DP KO HaCaT cells stably expressing DPI, DPIa, or DPII with a C-terminal mEGFP tag. Our results show the DP head domain position in the desmosomal plaque is isoform independent and the DP tail domain position correlates with rod length. The tail domain of DPI, the isoform with the longest rod, is furthest from the plasma membrane and that of DPII, the isoform with the shortest rod, is closest. We propose a variable tail location model to describe the architectural arrangement of each isoform. In this model, the DP isoforms are arranged with their rod domains parallel at an angle between 21° to 25° from the plasma membrane. These results provide valuable insight into the role of DP isoforms in desmosomal architecture and function.

## Introduction

Desmosomes are protein complexes that mediate cell-cell adhesion and provide integrity to tissues that experience mechanical stress (Kurn and Daly, 2024, Johnson et al., 2014, Bharathan et al., 2024). Dysregulation of desmosomes contributes to a variety of human diseases that primarily impact the skin and heart (Yuan et al., 2021, Mohammed and Chidgey, 2021, Stahley and Kowalczyk, 2015). Desmosomes are composed of desmosomal cadherins desmogleins (DSG) and desmocollins (DSC), armadillo protein family members plakophilin (PKP) and plakoglobin (PG), and desmoplakin (DP) (Fig 1A) (Kowalczyk et al., 1994, Kowalczyk and Green, 2013). Electron microscopy has revealed that desmosomes are organized into three distinct bands or regions, each oriented parallel to the membrane (Bharathan et al., 2024, Odland, 1958). In the extracellular space between two neighboring cells, the cadherin domains of DSG and DSC undergo trans binding and constitute the extracellular domain (ECD). The DSG and DSC cytoplasmic tails, PG, PKP, and the head domain of DP localize to the outer dense plaque (ODP). Finally, the DP tail domain localizes to the inner dense plaque (IDP) (North et al., 1999, Stahley et al., 2016). The specific 3D arrangement of proteins within these 3 domains remains unresolved.

**Figure 1:**
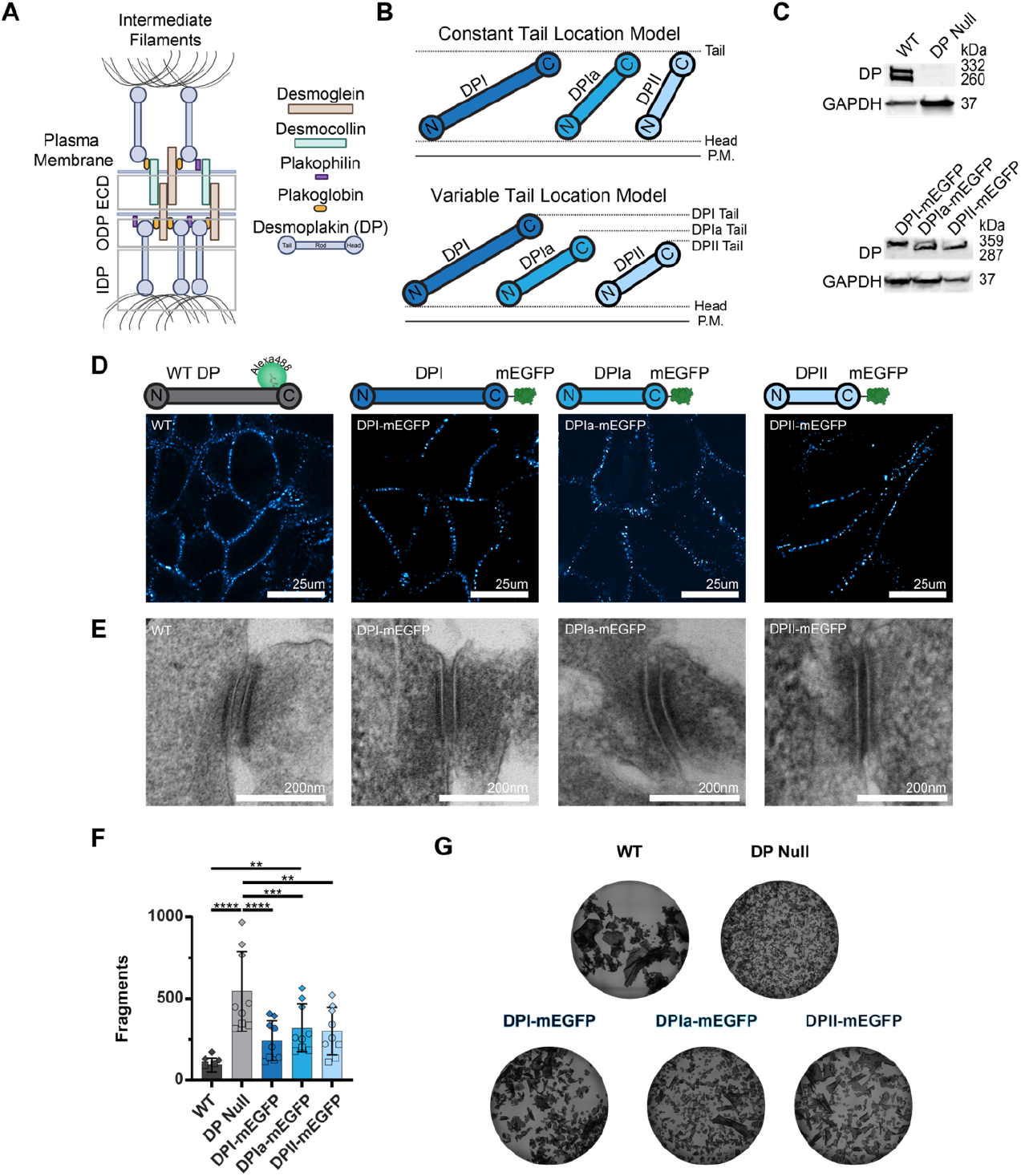
Characterization of Desmoplakin Isoform Expressing HaCaT Cells. **A)** Schematic of a desmosome, subunits, and domains. **B**) Two possible models of desmoplakin architectural arrangement: the constant tail location model (top) and the variable tail location model (bottom). **C**) Western blots showing DP expression in WT and DP KO HaCaT cells (top) and DP-mEGFP isoform expression in the DPI-, DPIa-, and DPII-mEGFP stable cells (bottom). **D**) Representative confocal images of DP in WT HaCaT cells and DP-mEGFP stable cells (scale bar = 25 μm). **E**) Representative TEM images of desmosomes in WT and DP-mEGFP isoform expressing HaCaT cells (scale bar = 200 nm). **F)** Number of fragments from WT, DP KO, and DP-mEGFP isoform HaCaT cells in a dispase cell adhesion assay (Mean +/-s.d., 3 replicates) WT HaCaT: 94 ± 43, DP null HaCaT: 544 ± 244, DPI-mEGFP: 242 ± 121, DPIa-mEGFP: 313 ± 155, DPII-mEGFP: 321 ± 123. One-way ANOVA: ****P<0.0001, **P<0.01, *P<0.05. **G)** Representative images from the dispase cell adhesion assay.

DP is an obligate desmosomal protein that is responsible for anchoring desmosomes to intermediate filaments (IFs), such as keratin. DP has a tripartite structure with a N-terminal plakin head domain, coiled-coil central rod domain, and IF-binding C-terminal tail domain. The head domain contains six spectrin repeats and an N-terminal unstructured region which interacts with PG and PKP (Bornslaeger et al., 2001, Kowalczyk et al., 1999). DP forms homodimers through the coiled-coil α-helical rod domain, and mutations destabilizing DP dimerization impair desmosome integrity (Daday et al., 2019, Kowalczyk et al., 1994, Green et al., 1992). The tail domain contains three plakin repeat domains, a linker domain, and a glycine-serine-arginine-rich region that modulates DP-IF interactions (Choi et al., 2002).

Alternative splicing of *DSP* generates three isoforms (DPI, DPIa, and DPII) that are structurally identical except for differences in the length of the rod domain (Cabral et al., 2010, Green et al., 1988). DPI (332 kDa) has a 888 AA rod domain, DPIa (279 kDa) has a 448 AA rod domain that is roughly half the length of the DPI rod, and DPII (260 kDa) has a 290 AA rod domain that is approximately one third of the DPI rod (Cabral et al., 2010, O’Keefe et al., 1989). The tissue expression of the isoforms is variable with stratified epithelia expressing DPI and DPII at roughly equivalent levels, while cardiomyocytes primarily express DPI. DPIa is a minor isoform with significantly lower expression (Cabral et al., 2010). The essential role of tissue specific expression of the DP isoforms is highlighted by rare human diseases. The loss of DPI expression by an isoform specific homozygous nonsense mutation leads to early onset cardiomyopathy (Uzumcu et al., 2006). Additionally, heterozygous premature stop codon variants generating either DPI or DPII haploinsufficiency can result in palmoplantar keratoderma involving skin thickening on palms and soles (Armstrong et al., 1999, Whittock et al., 1999). Knock down of DPI or DPII had differential effects on desmosome cadherin composition and mechanical resistance (Cabral et al., 2012). These findings suggest that the expression of multiple DP isoforms in skin provides greater resistance to mechanical strain, and the isoforms cannot substitute for one another in the heart. However, how DP isoforms individually contribute to desmosome function is unknown.

Desmosomes are complex, macromolecular assemblies that are essential in maintaining tissue integrity. While the identity of desmosomal components is known, their architecture is not well defined. The arrangement of desmosomal cadherins in the ECD has been determined to have repeating antiparallel architecture, but with a degree of flexibility (Dean et al., 2024, Sikora et al., 2020). Electron tomography has identified a repeating pattern of protein densities in the ODP, relating to PKP, PG, and the DP head domain (Al-Amoudi et al., 2011). Unlike other desmosomal components, DP spans between the ODP and IDP. DP-keratin binding has been defined, but this interaction within the IDP has yet to be characterized (Bartle et al., 2020, Kroger et al., 2013). DP head and tail domain plaque positions have been mapped through immunogold EM and direct stochastic optical reconstruction microscopy (dSTORM) (North et al., 1999, Stahley et al., 2016). Interestingly, this imaging data combined with *in vitro* measurement of DP head to tail “length” suggest the long axis of DP is oriented not perpendicular but at an acute angle relative to the plasma membrane (DP angle of alignment) (O’Keefe et al., 1989, Stahley et al., 2016). We have found that DP tail domain localization correlates with function during desmosome maturation and with ODP protein composition (Beggs et al., 2021, Stahley et al., 2016, Beggs et al., 2022). However, none of these studies compared DP isoforms and, in many cases, multiple isoforms were indistinguishable. Given the importance of DP isoform expression to desmosome function, deciphering their architectural arrangement is essential to understand the junction’s role in combating mechanical strain. In this study, we utilize human keratinocyte HaCaT cells expressing single DP isoforms and super-resolution dSTORM to uncover isoform dependent DP architecture.

## Results and Discussion

### Characterization of HaCaT Cells Expressing DP-mEGFP Isoforms

DP rod domain length varies between isoforms, but whether and how this affects desmosome structure and function is not understood. We sought to determine how the isoform specific rod lengths impact DP architecture and propose two possible models (Fig 1B). The “uniform tail position” model hypothesizes the tail domain of each isoform is the same distance away from the plasma membrane, creating a single interface for IF binding. In contrast, the “variable tail location” model hypothesizes the tail domain position is dependent on the rod domain length of each isoform. This could arise from the DP isoforms adopting the same angle of alignment (θ) relative to the plasma membrane within the plaque. In both models, the location of the DP head domain within the plaque is consistent across all isoforms, based on their identical sequences.

A central challenge to studying DP isoform architecture is their sequence identity. Specific antibodies for the smaller isoforms, DPIa and DPII, are not available preventing their individual labeling in wild type cells which express multiple isoforms. To address this challenge, we stably expressed DPI, DPIa, or DPII with a C-terminal mEGFP tag in CRISPR/Cas9 engineered DP knockout (ko) HaCaT human keratinocytes (Wanuske et al., 2021). This generated three HaCaT cell lines each expressing only a single DP-mEGFP isoform. While DP was not detected in the ko HaCaTs, overexpression showed each isoform with mEGFP tag at their expected molecular weights (Fig. 1C). All DP-mEGFP constructs localized to cell borders, similar to antibody labeled WT HaCaT cells (Fig. 1D). To examine desmosome ultrastructure, we carried out transmission electron microscopy. While desmosomes are not present in the parental ko cells (Wanuske et al., 2021), we found desmosome ultrastructure was indistinguishable between WT and the DP-mEGFP isoform expressing cell lines (Fig. 1E). Finally, intercellular adhesion was investigated in mechanical dissociation assays. Expression of mEGFP tagged DP isoforms partially rescued resistance to mechanical strain, which is severely disrupted by DP ko (Fig 1F, G). Interestingly, there was no significant difference in resistance to mechanical strain between DP isoforms indicating each isoform has a similar capacity to resist mechanical strain. However, the cells expressing a single isoform resisted mechanical strain less effectively than WT HaCaTs. This likely arises from a difference in DP expression level or could suggest the presence of multiple DP isoforms increases desmosome resistance to mechanical strain (Cabral et al., 2012, Spindler et al., 2014). Thus, the expression of DP-mEGFP isoforms in DP ko HaCaTs recapitulates desmosome formation, localization, and ultrastructure.

### DP Isoform Molecular Maps

Next, we wanted to determine the architectural arrangement of the different DP isoforms. To do so, we conducted super-resolution dSTORM on the DP-mEGFP expressing cells labeled with anti-GFP nanobodies. In all cell lines, DP appeared as puncta in diffraction limited widefield microscopy. dSTORM revealed two individual plaques within these puncta, each belonging to one half of a desmosomal junction (Fig 2A) (Stahley et al., 2016). To extract architectural information from these images, we quantified the distance between the plaques for many individual desmosomes in each group. Desmosomes have mirror symmetry across the midline, and this “plaque-to-plaque” distance represents the average separation of mEGFP on the tail domain of DP across neighboring cells. The larger the plaque-to-plaque distance, the further away from the membrane the tagged domain is located. We found the plaque-to-plaque distances for DPI-mEGFP, DPIa-mEGFP, and DPII-mEGFP tail domains were 186 ± 26 nm, 143 ± 18 nm, and 118 ± 14 nm respectively (mean ± s.d. Fig 2B, C). This data shows the position of the DP tail domain in the plaque is isoform dependent. The tail domain position of DPI-mEGFP, the longest isoform, was the furthest from the midline and DPII-mEGFP, the shortest isoform, was the closest to the midline. The rod domain of DPIa-mEGFP, is intermediate in length, and correspondingly its tail domain was located between that of DPI and DPII. Interestingly the distance of the DP tail domain from the midline correlates with the varying length of DP isoform rod domains.

**Figure 2.**
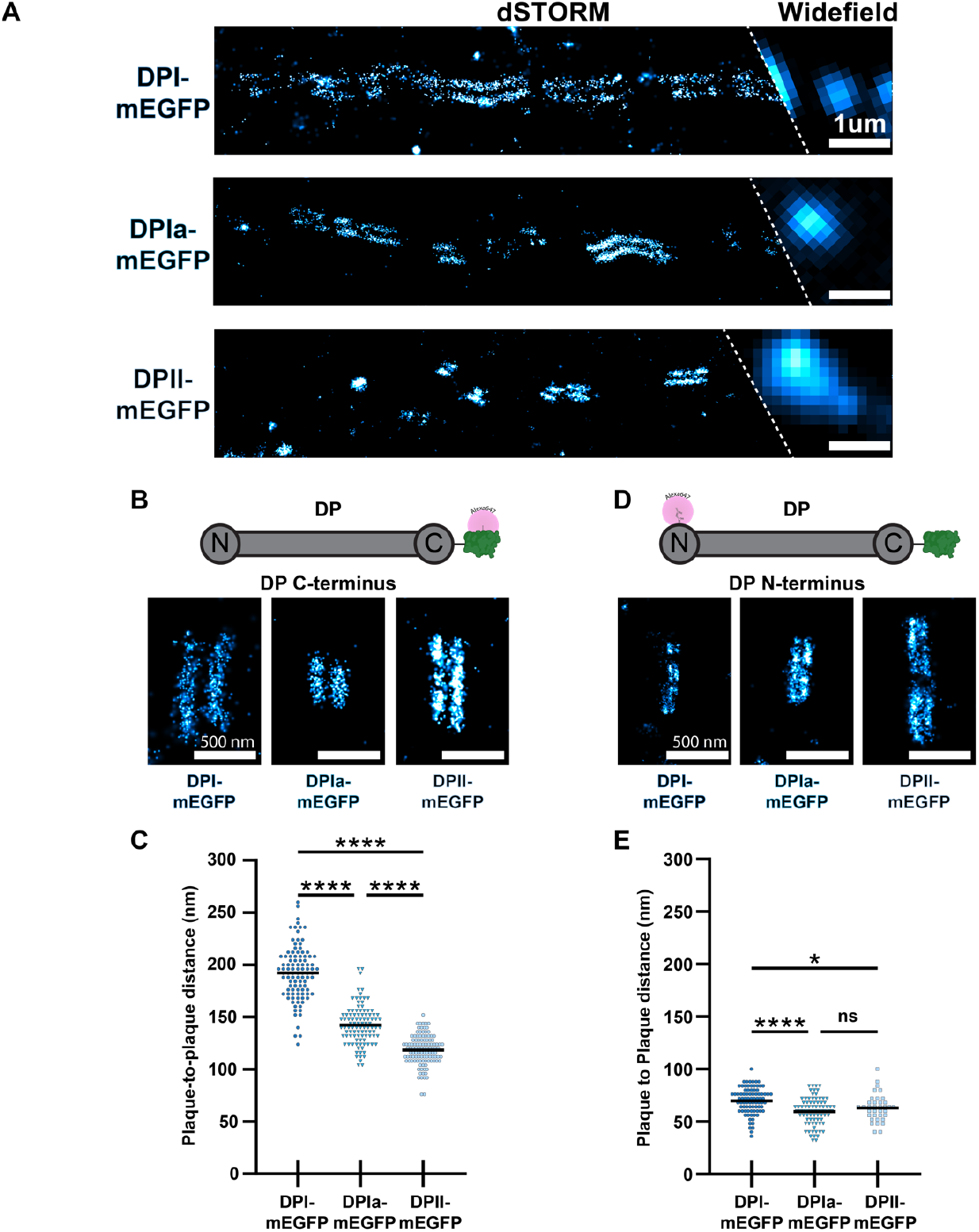
dSTORM Reveals the Architecture of Desmoplakin Isoforms. **A)** Representative widefield/dSTORM images of cells labeled with anti-GFP nanobody in DP-mEGFP isoform HaCaT cells (scale bar = 1 μm). **B**) Representative dSTORM images of tail domain of DP-mEGFP isoform HaCaT cells (scale bar = 500 nm). **C)** Scatter plot of the tail domain plaque-to-plaque distances for each isoform. DPI-mEGFP: N = 101, 192 ± 26 nm, DPIa-mEGFP: N = 101, 143 ± 18 nm, DPII-mEGFP: N = 112, 118 ± 14 nm. One way ANOVA with Tukey’s multiple comparison test: **** p < 0.0001. **D)** Representative dSTORM images of the head domain of DP labeled in DP-mEGFP isoform cells (scale bar = 500 nm). **E)** Scatter plot of head domain plaque-to-plaque distances for each isoform. DPI-mEGFP: N = 84, 70 ± 12 nm, DPIa-mEGFP: N = 79, 60 ± 13 nm, DPII-mEGFP: N = 60, 65 ± 15 nm. One way ANOVA with Tukey’s multiple comparison test: **** p < 0.0001 * p < 0.05.

Next, we tested if the differences in tail domain location arose from variation of the rod domain length, and not the overall position of DP. To do this, we quantified the position of the DP head domain by dSTORM following labeling with DP N-term antibody. The DP head domain plaque-to-plaque distance was smaller than that of the tail domain, reflecting its position in the ODP. The plaque-to-plaque distances were similar for all isoforms (DPI-mEGFP: 70 ± 12 nm, DPIa-mEGFP: 60 ± 13 nm, and DPII-mEGFP: 65 ± 15 nm) (Fig 2D, E). The relatively minor differences in the head domain localization are not sufficient to explain the 68 nm difference in the DPI and DPII tail domain plaque-to-plaque distances. Together, the similar position of the head domains with varying position of the tail domains supports the variable tail location model.

### DP Architecture is Isoform Dependent

To understand the precise arrangement of DP isoform architecture we combined this dSTORM data with published measurements of DP length. We determined the arrangement of DP within the plaque defined by the angle between the long axis of DP and the plane of the plasma membrane (θDP_x_). To determine the angle of alignment of DP, we utilized rotary shadow electron microscopy measurements which determined the average length of DPI to be 162 nm (O’Keefe et al., 1989). Because DPIa was not examined and DPII adopted a variety of conformations in measurements, we estimated their lengths. Using the measured head and tail domain lengths (16 nm each), we determined the DPI rod to be 130 nm long. Assuming rod length correlates with amino acid number and that the head and tail domains are conserved between isoforms; we calculated the expected lengths of DPIa (97 nm) and DPII (75 nm). With these predicted lengths and empirically measured head and tail domain positions for each isoform, we determined the DP angle of alignment to be 22°, 25°, and 21° for DPI, DPIa, and DPII respectively. These angles of alignment in coordination with the tail domain plaque-to-plaque distance measurement for each isoform strongly supports the variable tail location model (Fig 3A).

**Figure 3.**
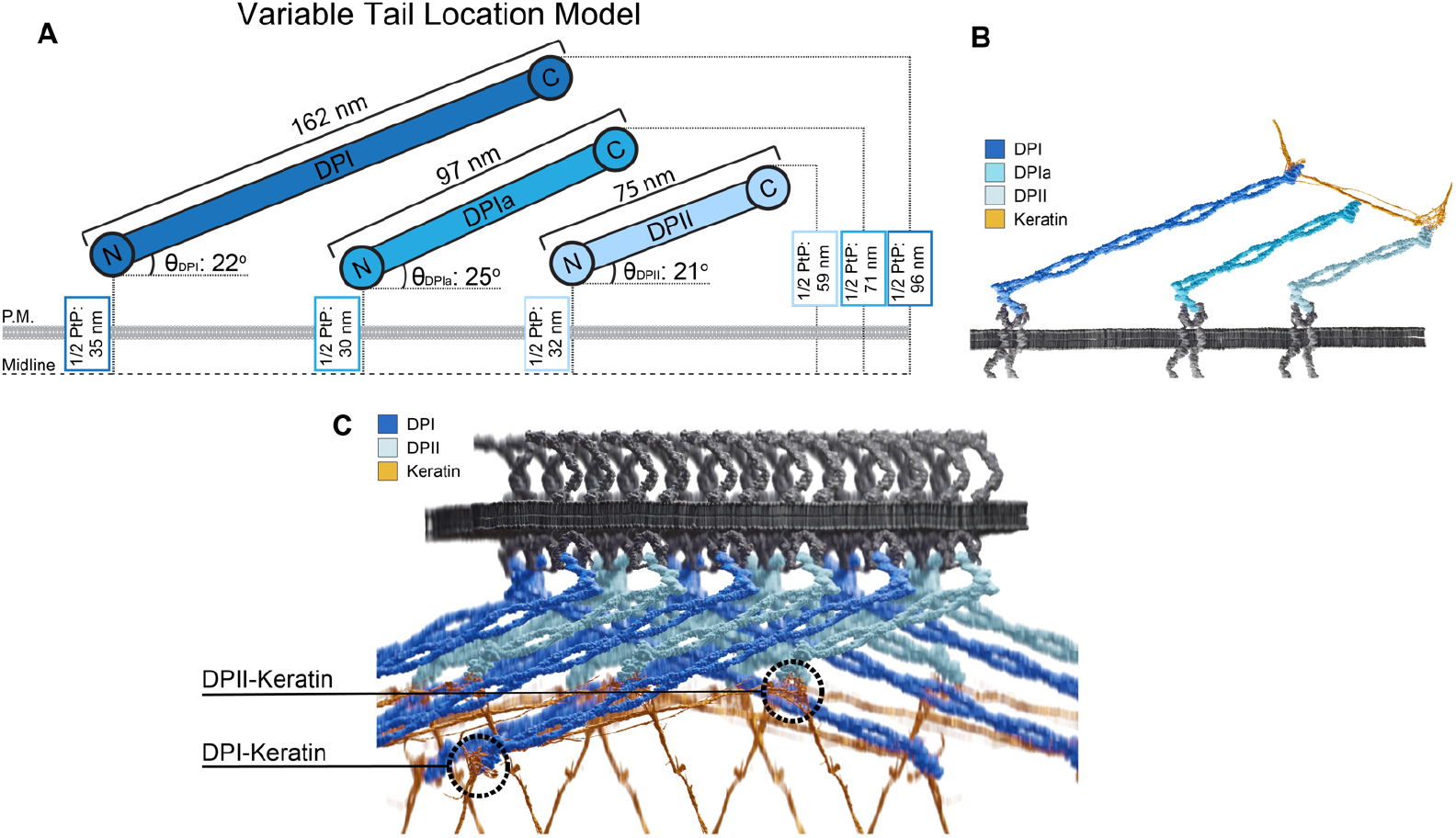
Desmoplakin Isoforms are Arranged in the Variable Tail Location Model. **A)** Scaled schematic of DPI, DPIa, and DPII architecture relative to the desmosome midline. 1/2 Plaque-to-plaque (1/2 PtP) distances were measured through our dSTORM analysis of each isoform. DP isoform lengths and angles of alignment were calculated as described in the text. There is cohesion in the predicted angle between the plane of the plasma membrane and the long axis of desmoplakin across the isoforms with θ_DPI_: 22°; θ_DPIa_, 25°, and θ_DPII_, 21°. **B)** Illustration of the variable tail location model with one keratin filament bundle binding to the tail domains of all three DP isoforms. **C)** 3-D model of a desmosomal plaque illustrating DPI and DPII isoforms with keratin looping through the plaque. Note the positions of DPI- and DPII-keratin interactions relative to the plasma membrane. Cadherins are arranged following anti-parallel model. Illustrations in 3B and 3C were created using Blender 4.0.

The head and tail domain plaque-to-plaque measurements for each isoform were acquired from desmosomes at the same stage of maturation. Despite the tail domain position varying between isoforms, the angles of alignment were within 4° of each other. As desmosomes mature, the composition of the ODP changes corresponding with a change to the position of the tail domain of DPI (Stahley et al., 2016, Beggs et al., 2022). We expect that a change to DP tail domain position would be reflected in all DP isoforms. While this would likely result in a new angle of alignment, we expect the angles would be similar amongst the isoforms.

Establishing the localization of the tail domain of DP isoforms generates exciting questions about the integration of keratin into the complex. The DP-keratin interaction is essential to both desmosome and keratin function. Post translational modifications to the keratin binding domain of DP directly influence desmosomal strength (Bartle et al., 2020, Hobbs and Green, 2012, Perl et al., 2023). Additionally, changes to keratin isoform expression impacts DP localization and desmosome strength (Wang et al., 2018). While the DP-keratin interface has been examined, how it varies between isoforms has yet to be described. The isoform-dependent tail domain position suggests that keratin could bind to DP at multiple layers relative to the plasma membrane. One scenario could involve one keratin bundle bound simultaneously by multiple DP isoforms at different levels as it loops though the IDP (Fig. 3B). In this model, multiple keratin-DP interaction interfaces might be present, either located distally from the membrane for DPI or at a relatively proximal position for DPII (Fig 3C). In skin epithelia, where DPI and DPII are expressed at relatively equal levels, such an interface could provide an advantage in resisting mechanical strain. It will be fascinating to study if greater functional integrity is supplied by multiple linkages and how variable DP rod domain lengths contributes to function. Overall, this work provides novel insights into the collaborative role of DP isoforms in the desmosome architecture.

## Methods

### Cell Culture

HaCaT cells were maintained in Dulbecco’s modified Eagle’s medium [DMEM; Corning, Tewksbury, MA] supplemented with 10% fetal bovine serum and 2% penicillin-streptomycin at 37°C and 5% CO_2_. DP KO HaCaT cells were a gift from the Spindler lab (Wanuske et al., 2021).

DP-mEGFP constructs were cloned to insert mEGFP following a linker (DPPVAT) on the C-terminus of each DP isoform. DPI-mEGFP, DPIa-mEGFP, or DPII-mEGFP was transfected into DP KO HaCaT cells using Lipofectamine 3000 following manufacturer instructions [Thermo Fisher Scientific, LC3000015]. Two days following transfection the GFP expressing population was enriched using fluorescence activated cell sorting (FACS) (BC FACSMelody). DP-mEGFP cells were maintained in media as described above supplemented with 500 ug/ml of geneticin.

### Antibodies

Antibodies used for immunofluorescence were: FluoTag-X4 anti-GFP conjugated with Alexa647 [N0304 NanoTag Biotechnologies, Gottingen, Germany]; FluoTag-X4 anti-GFP conjugated with Atto488 [N0304 NanoTag Biotechnologies, Gottingen, Germany]; anti-desmoplakin [A303-356A Bethyl Laboratries, Montgomery, TX]; anti-desmoplakin [00192 BiCell Scientific, Maryland Heights, MO]; anti-rabbit Alexa Fluor 647 [a-21244 Thermo Fisher, Waltham, MA]; anti-rabbit Alexa Fluor 488 [A-11008 Thermo Fisher, Waltham, MA]. Antibodies for western blotting were anti-desmoplakin [EPR4383(2) Abcam, Waltham, MA]; anti-GAPDH [6C5 Cell Signaling, Danvers, MA]; anti-mouse IgG HRP-Linked [7076 Cell Signaling, Danvers, MA]; and anti-rabbit IgG HRP-Linked [7074 Cell Signaling, Danvers, MA].

### Immunofluorescence

Cell fixation and labeling was conducted as described in Dean et al., (2024).

### Microscopy

Confocal images were obtained on a Ti-2 AXR microscope [Nikon Instruments, Melville, NY] equipped with a 60x 1.42 NA oil immersion objective, and 488 nm laser with Nyquist sampling. Z-stacks were acquired and deconvolved with the Richardson-Lucy algorithm with 8 iterations.

dSTORM images were obtained on a Nikon Ti-2 microscope system [Nikon Instruments, Melville, NY] equipped with a 100x 1.49 NA oil immersion objective, 647 nm laser, and Andor iXon EMCCD camera. 10,000 frames were acquired for each image. Samples were imaged in a 50mM Tris-HCl, 10mM NaCl, 10% glucose buffer with 5% 1M MEA [Sigma, St. Louis, Missouri] and 2% GLOX (20% 17mg/ml catalase [Roche, Penzberg, Germany] and 14 mg glucose oxidase [Sigma, St. Louis, Missouri]) each prepared in 50mM Tris-HCl and 10mM NaCl.

Protocols for TEM was conducted as described in Bartle et al., (2020). TEM was performed on a JOEL 1400 HC Flash TEM at 120 kV with an AMT NanoSprint43 Mk-II camera.

### Image Analysis

dSTORM images were analyzed as described in Beggs et al., (2021).

### Western Blot

Cells were grown to confluency, washed with ice cold PBS supplemented with a protease inhibitor cocktail [Roche Diagnostics GmbH, Germany], and scraped on ice. Samples were collected and centrifuged at 2,300 RPM at 4°C (5 m). The supernatant was aspirated, pellet was resuspended and lysed in 8 M urea on ice (30 m). Samples were centrifuged at 12,000 RPM at 4°C (15 m). The supernatant was collected and stored at -80°C. A BCA was utilized to determine protein concentration and either 85 or 15 ug of protein was loaded in 4-15% gradient gels for gel electrophoresis. 15 ug was utilized to confirm no DP was detectable in DP KO HaCaT cells while 85 ug was used to confirm expression of DP-mEGFP constructs. The samples were transferred to a PVDF membrane overnight at 4°C. The PVDF membranes incubated in Intercept Blocking Buffer [LiCor, Lincoln, NE] at RT (1 h), incubated in primary antibody diluted in Intercept Antibody Diluent [LiCor, Lincoln, NE] overnight at 4°C, washed 3x with PBST, and incubated in secondary antibody diluted int Intercept Antibody Diluent at RT (1 h), washed 3x with PBST, and imaged and analyzed on Bio-Rad ChemiDoc MP imaging system.

### Dispase Fragmentation Assay

Protocol was conducted as described in Dean et al., (2024). Three technical replicates with three biological replicates were obtained.

## Acknowledgements

The authors thank the High Resolution Imaging Facility at the University of Alabama in Birmingham for the excellent support and assistance with dSTORM and TEM.

## Author Contributions

Conceptualization: C.A., K.P., A.M.; Methodology: C.A., K.P., V.S., A.M.; Validation: C.A., K.P.; Formal Analysis: C.A., K.P.; Investigation: C.A., K.P., Y.T.; Resources: A.M., V.S.; Writing-Original Draft: C.A., K.P.; Writing-Review and Editing: C.A., K.P., Y.T., N.B., V.S., A.M.; Visualization: C.A., K.P., N.B., A.M.; Supervision: A.M.; Funding Acquisition: A.M.

## Funding

This work was supported by the National Institute of Health (NIH) National Institute of Arthritis and Musculoskeletal and Skin Diseases (NIAMS) R01 AR072697 to ALM and the UAB High Resolution Imaging Facility was supported by NIH National Cancer Institute (NCI) P30 CA013148.

## Competing Interests

The authors declare no competing or financial interests.

## Notes

### Competing Interest Statement

The authors have declared no competing interest.

